# Direct determination of oligomeric organization of integral membrane proteins and lipids from intact customizable bilayer

**DOI:** 10.1101/2023.03.12.532274

**Authors:** Aniruddha Panda, Fabian Giska, Anna L. Duncan, Alexander J. Welch, Caroline Brown, Rachel McAllister, Hariharan Parameswaran, Jean N. D. Goder, Jeff Coleman, Sathish Ramakrishnan, Frédéric Pincet, Lan Guan, Shyam Krishnakumar, James E. Rothman, Kallol Gupta

## Abstract

Hierarchical organization of integral membrane proteins (IMP) and lipids at the membrane is essential for regulating myriad downstream signaling. A quantitative understanding of these processes requires both detections of oligomeric organization of IMPs and lipids directly from intact membranes and determination of key membrane components/properties that regulate them. Addressing this, we have developed a platform that enables native mass spectrometry (nMS) analysis of IMP-lipid complexes directly from intact and customizable lipid membranes. Both the lipid composition and membrane properties (such as curvature, tension, fluidity) of these bilayers can be precisely customized to a target membrane. Subsequent direct nMS analysis of these intact proteo-lipid vesicles can yield the oligomeric states of the embedded IMPs, identify bound lipids, and determine the membrane properties that can regulate the observed IMP-lipid organization. Applying this, we show how lipid binding regulates neurotransmitter release and how membrane composition regulates the functional oligomeric state of a transporter.

## INTRODUCTION

The cellular membrane offers a chemically diverse environment that regulate oligomeric assemblies of the embedded integral membrane proteins (IMPs)^1–4^. A quantitative understanding of these oligomeric organization, and how they are regulated by the resident membrane, demands tools with high molecular resolution to detect and identify the assemblies of IMPs and lipids directly from intact and tunable membranes. The membrane tunability would further enable explicit changes in individual membrane constituents/properties, affording quantitative determination of their role in regulating the observed assemblies of the embedded IMPs. nMS, with its MS/MS capabilities, offers an unmatched molecular resolution to determine the oligomeric assemblies of proteins and their bound ligands^5–10^. Seminal works have also established nMS as a key technique to study IMPs^11–16^. However, nMS studies of IMPs requires prior in-solution dissolution of the intact membranes, typically by non-lipid bilayer forming membrane mimetics, such as detergents, bicelles, amphipoles, or mechanical disruption^11,13,17–20^. This points to a critical limitation, as these prior in-solution bilayer disruptions may fail to maintain essential properties of the native bilayer, such as the membrane curvature, fluidity, membrane tensions, and lateral mobility, which may regulate the physiological oligomeric states of IMPs^1,2,21^. While MSP-based nanodiscs do provide a bilayer environment^15,21,22^, their flat disc-like architecture provide limited control over the key membrane properties mentioned above.

In contrast, liposomes offer the most flexible in vitro bilayer platform^21^. Varying the composition of lipids, size, and protein-to-lipid ratio, vesicles can be precisely tuned to emulate the biophysical properties of a target physiological membrane^23,24^. Further, the lateral mobility offered by liposomes allows embedded transmembrane domains to interact with each other and with specific lipid domains, in a diffusion-controlled manner, as they do in the native membrane^23^. Consequently, liposomes are the most reliable in vitro platform for studying the functional activities of IMPs^25,26^. Nevertheless, previous attempts to use them as a carrier in nMS have only been successful to study soluble peripherally associated small peptides/proteins ^27,28^. To date, it has not been possible to study any IMPs directly from intact liposomes.

## RESULTS

To overcome this, we first established two independent pipelines that deliver nMS-ready lipid vesicles, embedded with the desired IMPs (Extended Data Fig. 1). Leveraging the flexibility of liposomes, the protocol enables us to mimic a target physiological membrane by tuning the lipid composition, curvature (by changing diameter), and fluidity (by changing the composition of cholesterol or unsaturated lipids), and the protein-to-lipid ratio of the vesicle.

Next, we developed a protocol that enables analysis of the embedded IMP-lipid complexes by directly subjecting these vesicles to nMS. Previous attempts to detect IMP from intact liposomes solely depended on collision induced activation (CID). This imposes a critical limitation, as CID energies available within commercial mass spectrometers are typically insufficient to break intact lipid vesicles and detect embedded IMPs. Indeed, previous work has demonstrated the necessity for prior in-solution dissociation of vesicles via mechanical disruption such as sonication or mild detergents to supplement CID activation^19,20^. Further, post-quadrupole collisional ejection of protein-protein/protein-lipid complexes from the bilayer prohibits downstream top-down MS/MS analysis of these complexes for unambiguous identification of bound lipids/proteins. Here, we report a multi-prong strategy to overcome these obstacles, allowing for the first time both MS and MS/MS analysis of IMP-lipid complexes directly from intact lipid vesicles.

First, to reduce the reliance on collisional activation alone, we exploited a unique surface biophysical characteristic of the bilayer. Our inspiration comes from previous works, mostly in the field of nanoparticle-based drug delivery, demonstrating how modulation of charges can aid the destabilization of lipid-bilayer^29^. Electrospray supercharging agents have been shown to manipulate surface charges of analytes, including MSP-peptide belts in the gas phase^22,30,31^. We hypothesized that similar surface charge manipulation of lipid bilayers by super-charging agents during the electrospray process may reduce the energy needed to ablate out and detect IMPs out of the bilayer. Next, we use the in-source trapping available in our MS platform to complement the collision cell CID. The current study uses Orbitrap Q-Exactive UHMR platform that offers conventional beam-type collision-cell CID (termed as HCD), as well as in-source trapping and activation at the inject flatapole where ions can be trapped and activated by creating a potential energy well between the S-lens-inject flatapole-inject flatapole lens^32^. Here, by modulating the negative potential gradient between the S-lens and inject flatapole, termed as ‘Desolvation energy’ in the instrument, the trapped ions can be collisionally activated through their collision with the neutral gas^32^. Here, we make use of this in-source trapping and activation to supplement the beam-type CID activation^32^. Together, beyond just supplementing the collision energy at the collision cells, these strategies can enable ablation of IMPs from the bilayer and allow MS and MS/MS analysis of protein-protein/lipid complexes directly from the bilayer.

To establish the broad applicability of our platform we chose a diverse set of IMPs of both prokaryotic and eukaryotic origin, with oligomeric states ranging from monomer to pentamer, masses 12kDa-226kDa, number of transmembrane-helices 1-28 (Table 1), and functions ranging from ion transport, water transport, sugar transport, mechanosensation to membrane fusion. To further illustrate the general applicability of the platform to any target physiological membrane, we reconstituted these proteins in a diverse set of liposomes that mimic the lipid composition of eukaryotic endoplasmic reticulum (ER), plasma membrane (PM), mitochondria, Golgi, and bacterial inner membranes ( Table 2)^33–35^. We confirmed the presence of vesicles in each of these samples through TEM imaging prior to nMS (Extended Data Fig. 2). Subsequent direct nMS analysis of each of these vesicles led to the detection of the embedded IMPs in their expected physiological oligomeric states (Fig. 1a, Extended Data Fig. 3). Table 1 shows excellent agreement between the theoretical and experimental masses and oligomeric states. Going beyond synthetic lipids, the platform also enables us to reconstitute proteins in lipids directly obtained from the host organism. Fig. 1b shows the spectra of two *E.coli* proteins, AqpZ and LacY, directly from liposomes that are prepared by using lipids directly extracted from *E Coli*, made up of 54 native lipids from all lipid classes (Fig. 1c and Extended Data Fig. 4). Hence, these vesicles truly capture the lipid diversity of the native-bilayer where AqpZ and LacY reside. The observed masses and oligomeric states are again in complete agreement with the theoretical values, confirming the ability of the platform to study proteins directly from the host-derived native lipid bilayer. Another major advantage of liposomes is their flexibility to make membranes of different membrane curvatures and fluidity. Leveraging this, we prepared proteo-vesicles of varying curvature by controlling the size of the liposomes (See Methods and Extended Data Fig. 2f,g,and h). This was confirmed through EM images of the liposomes before nMS analysis (Extended Data Fig. 2f,g,and h). To demonstrate that we can also successfully ablate IMPs from liposomes of different curvature, we reconstituted VAMP2 in each of these liposomes. Fig. 1d shows that for each of these cases, the embedded VAMP2 can be successfully detected from the liposomes of different curvature. While VAMP2 is not a curvature sensing protein (hence all three liposomes show identical spectra), the success paves the way to study effect of membrane curvatures in curvature sensing IMPS from liposomes of defined curvature.

**Table 1:**
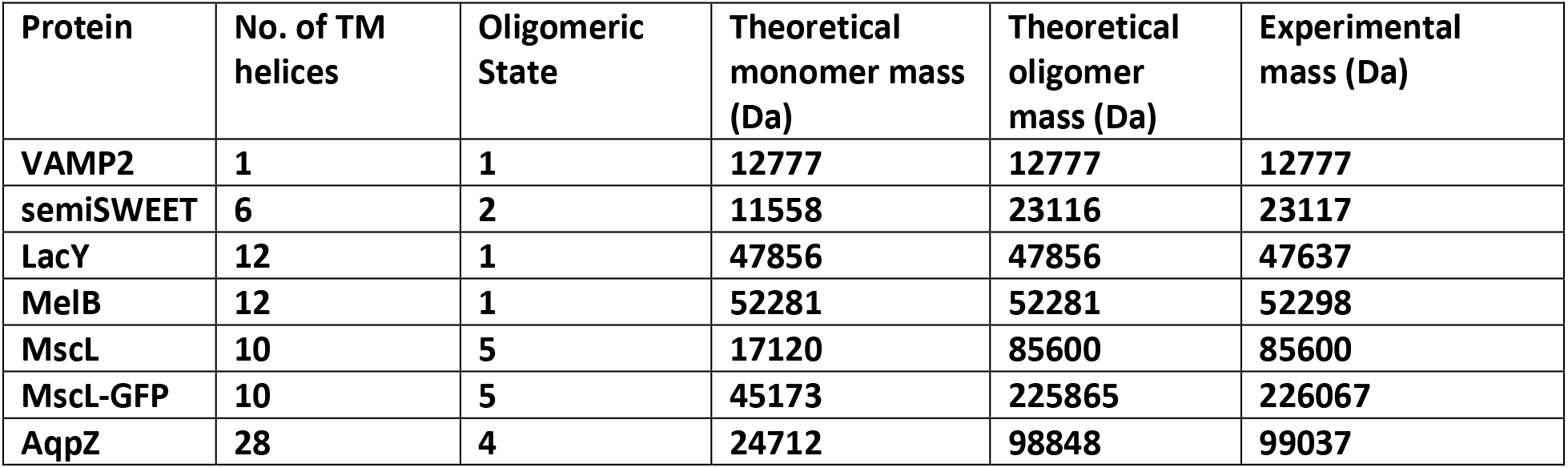
List of proteins used in this study with their theoretical and experimental masses, number of transmembrane helices, and oligomeric state

**Table 2:** Molar percentage of different lipids in different organellar membrane mimicking vesicles

| Lipid | Exact mass (Da) | ER | Golgi | PM | Mitochondria | Bacterial PM | Synaptic Vesicle |
| --- | --- | --- | --- | --- | --- | --- | --- |
| PC(18:1(9Z)/18:1(9Z)) | 785.60 | 54 | 45 | 23 | 37 | 0 | 21.3 |
| PE(18:1(9Z)/18:1(9Z)) | 743.54 | 20 | 17 | 11 | 31 | 67 | 13.7 |
| Cholesterol | 386.35 | 5 | 7.2 | 34 | 0 | 0 | 44.0 |
| PI(16:0/18:1(9Z)) | 836.52 | 9 | 9 | 4 | 6 | 0 | 2.4 |
| DAG(16:0/18:1(9Z)) | 594.52 | 3 | 0 | 0 | 3 | 0 | 0 |
| PS(18:1(9Z)/18:1(9Z)) | 786.52 | 4 | 4 | 8 | 1 | 0 | 8.0 |
| SM(d18:1/18:0) | 730.59 | 4 | 12 | 17 | 2 | 0 | 5.6 |
| Brain PIP2 | 1096.38(Formula Weight) | 0 | 0 | 2 | 0 | 0 | 0 |
| CL(1'-[18:1(9Z)/18:1(9Z)],3'-[18:1(9Z)/18:1(9Z)]) | 1457.04 | 0 | 0 | 0 | 16 | 10 | 0 |
| PG(16:0/18:1(9Z)) | 748.50 | 0 | 4.8 | 0 | 3 | 23 | 0 |
| Galactosyl($\alpha$) Ceramide (d18:1/16:0) | 699.56 | 0 | 0 | 0 | 0 | 0 | 2.4 |
| Brain Ceramide | 565.95(Formula Weight) | 0 | 0 | 0 | 0 | 0 | 1.6 |
| Lissamine Rhodamine PE(18:1(9Z)/18:1(9Z)) | 1284.5 | 1 | 1 | 1 | 1 | 1 | 1 |

**Fig. 1:**
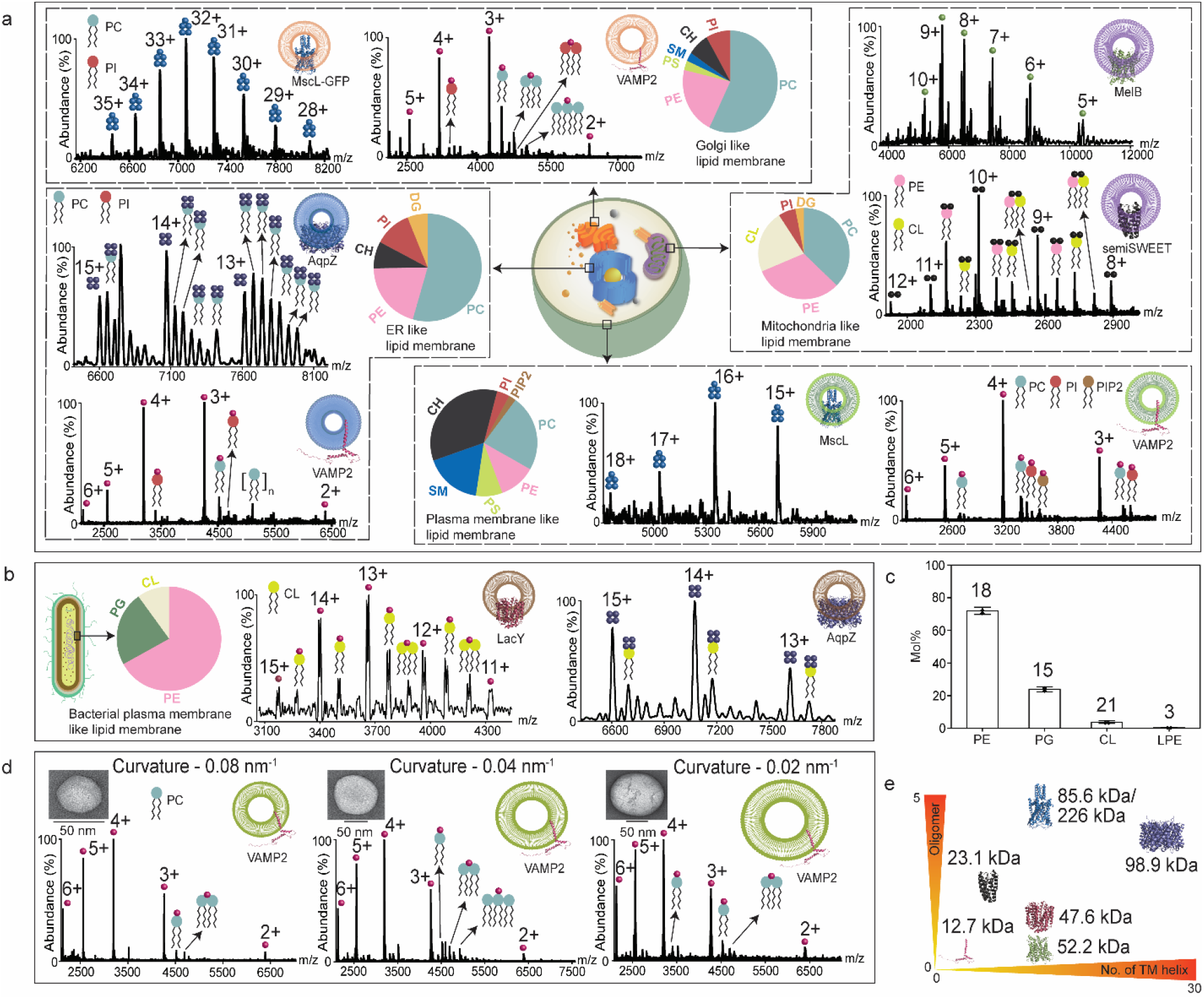
Detection of oligomeric organization of integral membrane proteins directly from bilayers mimicking physiological membranes. **a**, Oligomeric organizations of a range of IMPs detected directly from liposomes mimicking the bulk lipid composition of mammalian plasma membrane (PM), endoplasmic reticulum (ER), mitochondria (Mito), and Golgi membranes^33–35^. The lipid classes present and their relative composition in each of these membranes are depicted in the respective pi-charts and Table 2. These vesicles, reconstituted with the target IMPs were directly subjected to nMS. From each of the vesicles, the observed masses as well as the oligomeric states of the embedded IMPs are in complete agreement with the theoretical values of each of these proteins (Table 1). Besides observing the oligomeric states, the satellite peaks adjacent to the charge state of the apo-proteins also depicts the bound lipids adducts. The exact theoretical masses of the lipids used to form the bilayer are listed in Table 2. The identity of the bound lipids can be determined by directly comparing the mass of the adduct with these theoretical masses of the specific lipids used to form the bilayer ( Table 2, Extended Data Fig. 3) All measurements were replicated over three independent measurements. b, nMS spectra of AqpZ and LacY, two proteins that are endogenous to the *E.coli*, directly from bilayers consisted of native *E. coli* lipids. All measurements were replicated over three independent measurements. c, Lipidomics analysis revealed that these bilayers consist of more than 50 different lipids from three different lipid classes. The data is plotted as mean ± s.e. (N=2). d, nMs analysis of IMPs directly from bilayers of different vesicle size and curvature. The EM images shown in inset further confirms generation of vesicles of desired diameters. All measurements were replicated over three independent measurements. e, Shows the collection of standard IMPs chosen that broadly ranges masses from 12-226kDa, no of TM helices 1 - 28, and oligomeric states 1-5.

Next, we made a series of liposomes with decreasing membrane fluidity by increasing the cholesterol (CH) percentage. We confirmed the decrease in membrane fluidity with fluorescence recovery after photobleaching (FRAP) experiments (Extended Data Fig. 5). Again, our data clearly shows that we can successfully detect the embedded protein from each of these vesicles.

Beyond detecting the target protein, we also observed lipid-bound species, elucidating which of the bilayer lipids specifically bind to the embedded proteins. The identity of the lipids can be determined by directly comparing the mass of the adduct with the theoretical masses of the lipids used for making bilayer ( Table 2, Extended Data Fig. 3). However, in cases where disparate lipids are close in mass, distinguishing them through the measured mass of the protein-lipid complex can be challenging. Alternatively, our ability to ablate the IMPs from the bilayer before the quadrupole enables us to circumvent this problem by isolating the lipid-bound species in the quadrupole and performing high-resolution MS/MS experiments on the lipid-protein complex (Discussed later, See Fig. 2a).

**Fig. 2:**
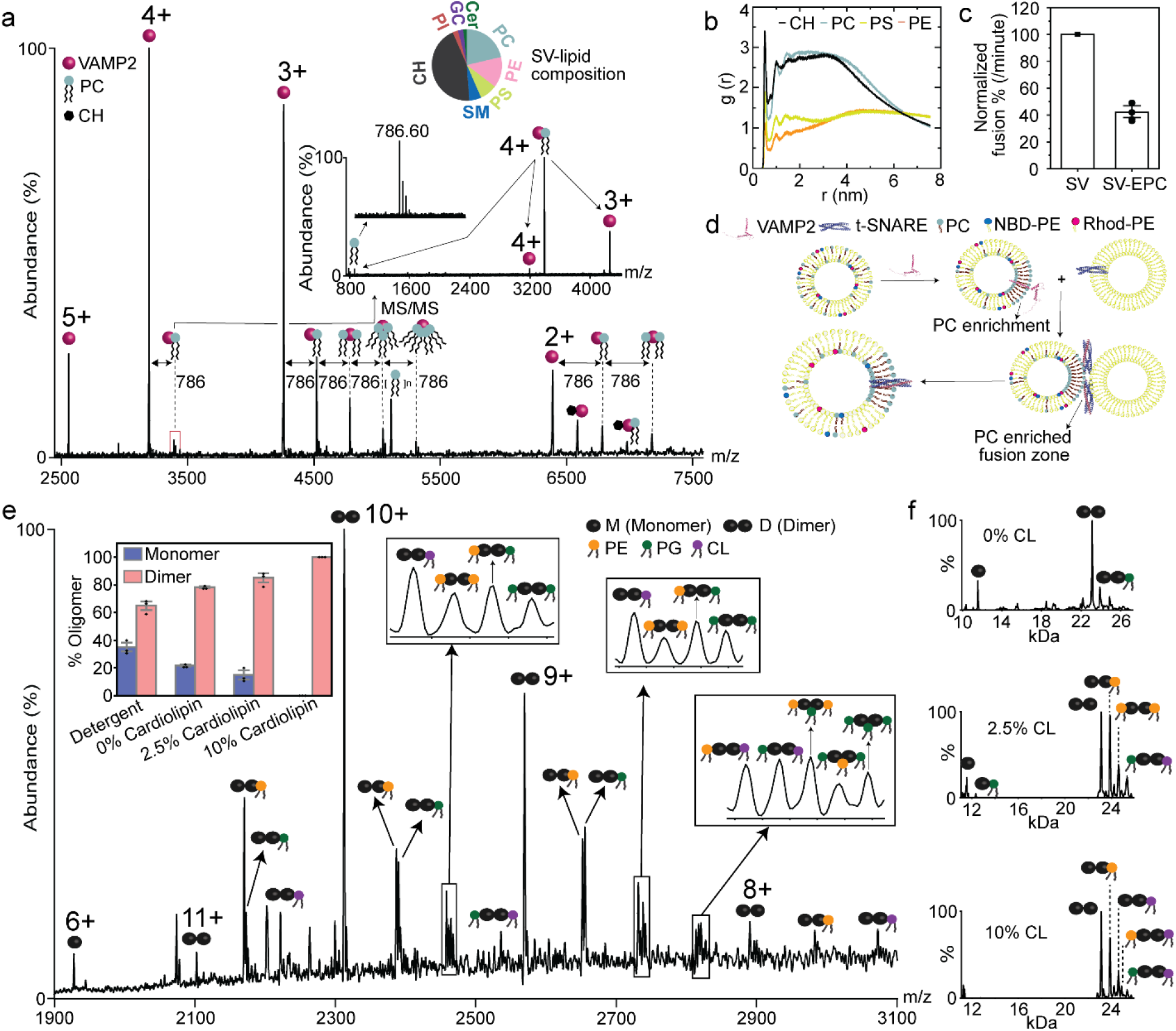
Direct analysis of VAMP2 and semiSWEET from native-like membrane. **a**, nMS of VAMP2 directly from vesicles that mimics the lipid composition, diameter, and curvature of SV. Each charge state of apo-VAMP2 was accompanied by a series of successive 786Da mass addition. To identify the adduct, we isolated the lipid bound 4+ charge state, subjected it to HCD MS/MS, and recorded a high-resolution spectrum (100,000). The m/z of the ejected lipid (786.60) matched exactly with the theoretical m/z of DOPC (786.60, error<5ppm), confirming the series of 786Da adducts observed for all charge states of VAMP2 to be successive PC binding. All measurements were replicated over three independent measurements. b, The radial density of different phospholipids around the juxta-membrane region of VAMP2, obtained through Coarse-grained MD simulation of VAMP2 in lipid bilayer. The data shows clear preference of PC and CH near the juxta-membrane region. c, Vesicle fusion assay performed between t-SNARE and VAMP2, which are reconstituted in vesicles with and without PC. For the later, PC was replaced by ethyl-PC (EPC). For both the cases, VAMP2 vesicles contained 1% Rhod-PE and NBD-PE. For each replicate the rate of fusion of EPC-SV vesicles are normalized to the corresponding control fusion rate from SV-like vesicles, taking the later as 100%. The data shows close to 60% reduction in vesicle fusion rate upon abrogation of PC binding to VAMP2. The data presented as mean ± s.e. (N=5). d, Schematic showing VAMP2-induced PC clustering ensures the segment of the SV-bilayer fusing with t-SNARE bilayer is enriched in PC, which may in turn reduces the energy of activation of fusion and speeds up the fusion rate. e, nMS of semiSWEET from a bilayer mimicking inner membrane of gram-negative bacteria showing almost exclusive presence of dimeric semiSWEET (>95%). Inset shows the monomer-dimer % as a function of CL% in the membrane. The data shows increasing CL% corresponds with increasing dimeric population in the bilayer. All measurements were replicated over three independent measurements. f, Mass plot obtained from UniDec analysis of semiSWEET from bilayer with 0%, 2.5%, and 10% CL. All measurements were replicated over three independent measurements.

Together, this data highlights the broad applicability of our platform to study IMPs directly from bilayers that can be customized to mimic the lipid composition, curvature, and fluidity of a target physiological membrane. Further, we demonstrate the ability to study IMPs directly from bilayers formed by the native host lipids. An interesting observation was as long as the liposomes were unilamellar, we did not find supercharging to be a universal requirement to ablate IMPs out of bilayer, with some proteins such as MscL, VAMP2 not requiring any supercharging (See Methods). This may be due to the inherent thermodynamic cost associated with ablating an IMP from bilayer, indicative of the solvation energy of the transmembrane domain in that bilayer. Future systematic studies using model transmembranes in different bilayer compositions can yield further mechanistic insight into this process.

### APPLICATION OF THE METHOD IN NEURONAL EXOCYTOSIS

Having established the broad applicability of our platform, we first focused on applying this to VAMP2, residing in synaptic vesicles (SV). In the neuronal synapses, neurotransmitter filled SVs dock and fuse with the plasma membrane (PM) and release neurotransmitters. The fusion of the SV with the PM takes place via complex formation between VAMP2, present in the SV, and the t-SNARE (a complex of SNAP25+Syntaxin) present at the PM^36^. An outstanding question is how the neuronal SNAREs attain their sub-millisecond time-scale of fusion. Higher-order molecular organization, including transmembrane-induced lipid-clustering, has been proposed to play a major role^37^. Using our approach, we sought to understand how VAMP2 can regulate lipid clustering at the SV membrane and thus regulate fusion. nMS analysis of VAMP2 from vesicles mimicking the SV in terms of the lipid composition, dimension, membrane fluidity, and membrane curvature^38^ led to direct detection of the protein (Fig. 2a). The apo-protein charge states were found to be accompanied by a series of 786Da adducts (Fig. 2a), which is close to the exact mass of both DOPC (785.59Da) and DOPS (786.53Da) and cannot be unambiguously resolved from MS1 alone. Here our ability to ablate VAMP2 from the bilayer just by front-end in-source trapping and activation enabled us to specifically quadrupole isolate the lipid-bound species (Δ = +786). During quadrupole isolation of the lipid bound VAMP2, we did not observe any leakage of lipids (Fig. 2a, Extended Data Fig. 6a). Subsequent high-resolution MS/MS analysis of the isolated lipid-protein complex (with Δ = +786) unambiguously yielded the identify the bound lipids as PC. A point to note that higher % of lipid bound VAMP2, compared to the apo-VAMP2, were observed for the lower charge states (Extended Data Fig. 6b). This is plausibly because higher charge states undergo a higher degree of unfolding in the gas phase, resulting in preferential loss of ligands^39,40^. But we chose the lipid bound species to the 4+ charge state for MS/MS as higher number of protons increases the possibility of observing a charged lipid loss. To systematically rule out the observed PC bindings as an artifact of head-group mediated non-specific polar interactions, we first repeated the experiments from liposomes where all PCs were replaced by sphingomyelin (SM), a lipid that has the same phosphocholine headgroup as PC but a different backbone. No detectable amount of SM binding was observed (Extended Data Fig. 7a). We then replaced all PCs in the bilayer with EPC, which has the same backbone as PC but an esterified phosphocholine headgroup. Again, no detectable amount of EPC binding was observed (Extended Data Fig. 7b). Together, these experiments confirm that the VAMP2-PC interaction is specific and requires both the glycerophosphate backbone and the headgroup of PC.

Additionally, we also observed binding of cholesterol (CH, 386.65Da). Notably, despite the established biochemical roles of CH in regulating the structures and functions of IMPs, successful MS-based detection CH binding has only been achieved by elegantly crosslinking CH to target IMPs^41^. To date CH binding has not been observed in nMS. This recalcitrance of IMP-CH complex to nMS is possibly due to the localization of CH in specific domains of eukaryotic membranes^1^. CH association with IMPs is often driven by localization of the proteins in those specific membrane domains, an environment that cannot be emulated in detergents and hence, had remained unobserved.

Surprisingly, even in our experiments, when detergent-solubilized VAMP2 was incubated with the same mixture of lipids used to form the SV-like liposome, no CH binding was observed (Extended Data Fig. 8). Further, the data clearly shows that VAMP2, in the detergent micelle, has no preferential binding towards any specific lipids, including PC. This clearly indicates that the observed PC-CH binding preference is specific to bilayers and is influenced by the SV-membrane environment. To rationalize this, we performed coarse-grained MD simulations of VAMP2 in a lipid-bilayer, resembling that of the SV membrane. The results (Fig. 2b), mirroring the experimental observations, clearly show that the juxtamembrane region of VAMP2 has a distinct preference to bind and associate with the PC-CH rich segment of the membrane. Indeed, previous works have established the distinct specificity of CH to partition with lipids with choline-headgroup, such as PC/SM, through H-bonding^1^. Our data indicate that in SV-membranes, VAMP2 localizes in a PC-CH enriched domain and imparts the observed lipid binding specificity. Hence, when studied from the detergent micelle, where such membrane domains cannot form, no such specificity towards PC-CH is observed. Another critical experimental evidence in support of this is the fact that selective removal of PC from the vesicles also abrogated CH binding (Extended Data Fig. 7). Together, this further emphasizes the critical need to study IMPs directly from bilayer. Intriguingly, previous studies have indicated a role of the juxtamembrane region of VAMP2 in regulating SV fusion, and the role of lipid binding was postulated as a possible rationale^42,43^. However, the identity of the bound lipids or their mechanism of action has remained absent. Among the two lipids we observed, CH is necessary for neuronal exocytosis^44^, with depletion of CH level shown to directly impair the release of secretory contents^45,46^. Our data provide direct evidence that VAMP2 directly binds to CH in the SV membrane and regulates the time-scale of vesicle fusion.

Further, our data show that VAMP2 binds to multiple PC and enriches the surrounding membrane with PC. This means that when an SV-resident VAMP2 forms a complex with a PM-resident t-SNARE, the segment of the SV bilayer around VAMP2 that faces and eventually fuses with the PM is enriched with PC (Fig. 2d). Intriguingly, previous work has shown that under the synaptic cleft conditions PC induces membrane fusion^47^. This suggests a mechanism in which VAMP2, through its specific lipid binding, enriches PC in the proximal SV-membrane and thereby reduces the activation energy for bilayer fusion at that membrane locus (Fig. 2d).

To test this hypothesis, we performed a fluorescence-based bulk fusion assay in the presence and absence of PC in SV-like vesicles (see Methods)^36^. In non-PC-containing vesicles we replaced PC with EPC, a PC analog that we showed does not bind to VAMP2 (Extended Data Fig. 7b). The fusion rates of both vesicles were independently measured by fusing them with PM-like vesicles containing t-SNAREs. Indeed, the rate of vesicle fusion decreased upon abrogation of PC binding to VAMP2 in SV-EPC vesicles (Fig. 2C, Extended Data Fig. 9). We separately confirmed that this decrease is not because of the introduction of EPC in the bilayer (Extended Data Fig. 9c). While the current work raises further questions that we are actively pursuing-such as where exactly in the juxtamembrane region PC binds, and how it regulates bilayer fusion, it demonstrates our platform’s ability to determine the lipid-binding specificity of a membrane protein. By directly studying IMPs from bilayers that mimic a target physiological membrane, and performing downstream MS and MS/MS, this method allows us to unambiguously determine how a particular IMP organizes the lipid-bilayer composition around itself through specific lipid-protein interactions.

### APPLICATION OF THE METHOD IN STUDYING LIPID INDUCED OLIGOMERIZATION OF MEMBRANE PROTEIN

We next sought to establish how the tunability of our platform allows us to determine the effect of independent bilayer constituents/properties on regulating the oligomeric organization of proteins in the bilayer. For this, we chose to examine a bacterial sugar transporter, semiSWEET. While previous detergent-based studies on semiSWEET have shown the presence of both monomeric and dimeric species, functional studies have pointed to an obligatory dimer^4^. To address this disparity, we first reconstituted semiSWEET into liposomes that mimic the lipid composition of the bacterial inner membrane. nMS analysis of these vesicles showed exclusively dimeric semiSWEET, establishing that semiSWEET in the bilayer is exclusively dimer (Fig. 2e). Next, we sought to find out if specific membrane constituents regulate this dimer formation. Previous studies have shown that cardiolipin (CL) co-purifies along with detergent extracted semiSWEET^4^. Considering these findings, we naturally gravitated towards studying the effect of CL first. To do this, we reconstituted semiSWEET in vesicles with an increasing amount of CL and compared the monomer-dimer distribution amongst these vesicles under the same MS conditions (Fig. 2e, Extended Data Fig. 10). As shown (Inset to Fig. 2e and inset to Extended Data Fig. 10a), the removal of CL from the bilayer led to a significant decrease in the dimer population and an increase in the monomer population. Correspondingly, an increase in CL% led to a concomitant increase in the dimer population, with physiological-like lipid composition showing an almost exclusively dimeric semiSWEET population. Interestingly, a recent MD simulation of semiSWEET on a bacterial inner membrane-like bilayer showed a higher abundance of CL in the oligomeric interface of the functional dimeric form^48^. Our data provide clear experimental evidence of how the presence of CL in the bilayer directly regulates this functional oligomeric state of the protein, and thus regulates sugar transport.

## DISCUSSION

Our current work presents a broadly applicable platform that can be used to study the molecular organization of proteins and lipids directly from bilayers of desired lipid composition and tailored biophysical properties. The tunability of this platform further allows us to delineate the role of independent membrane constituents and biophysical membrane properties in regulating observed protein assemblies. From GPCRs to ion channels, how specific lipid-binding and key membrane properties regulate higher-order protein-protein organization is a key unanswered question. Combining the unmatched ability of nMS to dissect protein-lipid assemblies with the tunability of customizable membranes will find broad applicability in studying these outstanding questions at molecular resolution.

## ACKNOWLEDGEMENTS

The authors acknowledge the late Ronald Kaback for several critical insights on making lipid bilayers. This work was partly supported by the National Institutes of Health grants R01GM141192 to KG, R01DK027044 to JER and SSK, and R01GM122759 and R21NS105863 to L.G. AJW was funded by a Pembroke College (University of Oxford) Rokos Award. ALD was funded by the Biochemistry Department, University of Oxford and Pembroke College Oxford.

## AUTHOR CONTRIBUTIONS STATEMENT

A.P. and K.G. designed the research and established the method. A.P. performed all experiments on VAMP2, semisweet, and MscL. F.G. performed the experiments on AqpZ, MelB, and LacY. A.D. and A.W. performed molecular dynamics simulations. C.B. performed the negative stain EM experiments and data analysis. R.M. performed lipidomics experiments and data analysis. H.P. and L.G. provided the purified MelB and LacY. J.N.D.G performed the FRAP experiments under the supervision of F.P. J.C. prepared the VAMP2 construct. A.P. performed the vesicle fusion experiments under the guidance of S.R. S.K. evaluated and provided critical feedback for bulk fusion experiments. J.E.R. provided supervision on all experiments on VAMP2 and synaptic vesicle and provided general critical insights. K.G. and A.P. wrote the manuscript with contributions from all other authors.

## CORRESPONDS

All corresponds should be addressed to

## METHODS

### RECOMBINANT PROTEIN EXPRESSION AND PURIFICATION

Recombinant v-SNARE (VAMP2) and t-SNARE (SNAP25+syntaxin1A) were expressed and purified as previously described^36,49,50^. Briefly, proteins were expressed in the *E. coli* BL21 strain using 0.5mM IPTG for 4 hours at 37°C. Cells were pelleted and lysed using a cell disruptor (Avestin, Ottawa, Canada) in HEPES buffer (25mM HEPES, 400mM KCl, 4% Triton X-100, 10% glycerol, pH7.4) containing 1mM DTT. Samples were clarified using a 45Ti rotor (Beckman Coulter, Atlanta, GA) at 142159xg for 30 minutes and subsequently incubated with Ni-NTA resin (Thermofisher Scientific, Waltham, MA) overnight at 4°C. For t-SNARE, the resin was washed with HEPES buffer supplemented with 50mM Imidazole, 1% octylglucoside (OG), 1mM DTT pH 7.4. Proteins were eluted using HEPES buffer supplemented with 500mM Imidazole, 1% octylglucoside (OG), 1mM DTT, pH 7.4. For v-SNARE resin was washed with HEPES buffer supplemented with 15mM Imidazole, 1% octylglucoside (OG), 1mM DTT, and then with HEPES buffer supplemented with 25mM Imidazole, 1% octylglucoside (OG), 1mM DTT pH 7.4. The resin was incubated with SUMO protease in HEPES buffer supplemented with 1% octylglucoside (OG), 1mM DTT overnight at 4°C. The next day, the protein was collected as flow-through from the gravity flow column.

semiSWEET, MscL, and AqpZ were purified as described previously^4,13^. Briefly, 20 ml of overnight Rosetta (DE3)pLysS cells with plasmids coding semisweet, MscL, and AqpZ were grown overnight. Then after refreshing the culture, bacteria were grown to OD – 0.8. At this point, the protein expression was induced with IPTG at a final concentration of 0.2 mM, and the bacteria were grown at 37°C for 4 hours except for semiSWEET (22°C for 15 hours). Cells were harvested by centrifugation (4000xg, 10 minutes, 4°C) and then resuspended in 100 ml of lysis buffer (20 mM Tris, 300 mM NaCl, pH 7.4 supplemented with protease inhibitor cocktail tablets (Pierce protease inhibitor mini tablets). Cells were lysed using a cell disruptor (Avestin, Ottawa, Canada) and then cell debris was removed by centrifugation (20,000xg, 20 minutes, 4°C). The supernatant was taken and the membranes were pelleted by ultracentrifugation (100,000xg, 2 hours 15 minutes, 4°C). The membranes were homogenized in 30 ml ice-cold membrane resuspension buffer (20 mM Tris, 100 mM NaCl, 20% Glycerol, pH 7.4) using a Potter-Elvehjem Teflon pestle and 2% (w/v) powder DDM was added except for MscL (1%OGNG was used) and left to tumble at 4°C for 2 hours. The samples were clarified by centrifugation (20,000xg, 40 minutes, 4°C) and then the supernatant was filtered through 0.22 μm filters. Then proteins were first purified by His-tag affinity chromatography in 20 mM Tris, 150 mM NaCl, 10% Glycerol, 0.02% DDM, pH 7.4. Then protein was cleaved from respective tags using appropriate protease (semiSWEET - Precission Protease, AqpZ, and MscL–TEV protease) and further purified using reverse Ni chromatography and size exclusion chromatography. During size exclusion chromatography the detergent was exchanged to OG for protein reconstitution into lipid vesicles.

LacY and MelB was purified as described previously^51,52^.

### PREPARATION OF NATIVE MS READY PROTEO-LIPID VESICLES

#### A. Density gradient float-up method

All lipids were purchased from Avanti Polar Lipids. For each of the vesicle preparation, the composition of the lipids is described in Table 2. The lipid mixtures were dried down in 12 x 75 mm Borosilicate glass tubes by a gentle stream of nitrogen, and any remaining traces of organic solvents were then removed under vacuum for 1 hour. A sample to produce protein-containing vesicles was prepared by dissolving the lipid film in 100μl of a reconstitution buffer containing the desired amount of the purified protein (lipid: protein ratio in the range of 100 to 1000 for different proteins). The lipids were dissolved in this solution by gentle agitation for 30 minutes at room temperature. Vesicles were then formed by rapid dilution by adding 200μl respective reconstitution buffer supplemented with 2mM DTT without detergent. Then detergent was removed by dialysis in constant flow dialyzing cassette using Spectrapore 6-8 kDa cutoff dialysis membrane against 4 liters of respective reconstitution buffer supplemented with 2mM DTT at 4°C. Vesicles were recovered and concentrated by floatation in a Nycodenz (Sigma) step gradient. Each 300μl dialysate was mixed with 300μl of 80% (w/v) Nycodenz dissolved in reconstitution buffer and then divided equally into two 5 x 41 mm ultracentrifuge tubes (Beckman). Then, each sample was overlayed with 300μl 30% (w/v) Nycodenz in reconstitution buffer followed by 100μl reconstitution buffer without Nycodenz. The samples were then centrifuged in an SW55 rotor (Beckman) at 279482xg for 4 hours at 4°C. The vesicles were carefully collected from the 0 −30% Nycodenz interface from each tube and combined. These samples were then analyzed for size in electron microscopy and dynamic light scattering (DynaPro Nanostar, Wyatt Technology). 30μl of these samples were dialyzed in constant flow dialyzing cassette using Spectrapore 6-8 kDa cutoff dialysis membrane against 4 liters of mass spec buffer (200 mM ammonium acetate, 2mM DTT) at 4°C. These dialyzed samples were used for nMS.

#### B. Sephadex G50 column method

This method was adapted from the previous work that enabled reconstitution of membrane proteins into liposomes for CryoEM structure determination of membrane protein directly from proteoliposomes^53^. The day before the MS experiments, Sephadex G-50 powder was dissolved in ammonium acetate buffer and sonicated in a water bath for 5 minutes. This suspension was then swelled overnight while degassing under a vacuum. On the day of the experiment, the Sephadex column was prepared by filling an empty column packed with the pre-swollen Sephadex gel. Separately, dried lipid film was resuspended in ammonium acetate buffer (200 mM ammonium acetate, 2 mM DTT). The lipid: protein ratio in the range of 100 to 1000 for different proteins. Then the solution was sonicated for 15 minutes in a bath sonicator and ten freeze-thaw cycles were performed (liquid nitrogen was used for freezing and a water bath set at 50°C was used for thawing). Then the appropriate detergent was added to the final concentration 2X CMC. This solution was then kept on ice for 30 minutes. After a 30-minute incubation, the desired amount of protein in 2X CMC detergent was added and the mixture was incubated on ice for 2 hours. This sample was placed on top of the prepared column and separated through Gel filtration to collect the fraction containing the proteoliposomes. All liposomes were prepared using 1% fluorescent lipid to conveniently track the elution of the liposomes. As a control, the same amount of protein in 2xCMC detergent was added to a blank buffer solution containing no liposomes and passed through sephadex G-50 column. Here, no proteins were observed in the elution volume corresponding to the elution volume of the proteoliposomes in the sample. Part of the proteoliposome samples were then analyzed using electron microscopy and dynamic light scattering (DynaPro Nanostar, Wyatt Technology) to confirm the size. If needed, an optional step of Nycodenz floatation, as described above, can be added to further separate the proteoliposomes from the empty liposomes.

#### C. Modulating curvature of nMS ready liposomes

Curvature of a curved surface is related to the radius of curvature of that point by the formula:

k = 2/(R); where k is the mean curvature and R is the radius of curvature.

For a spherical liposome this R translates to the radius of the liposome. Hence, by precisely varying the radius of liposomes the curvature of the membrane can be modulated. To create liposomes of specific radius, we extruded the liposomes (prepared either by Method A or B) through specific size membrane filters 21 times. The size of the filter defines the diameter of the liposomes. In the current work, for Fig. 1d, to generate liposomes of 0.08, 0.04, and 0.02 nm^-1^ curvatures, we extruded the liposomes through a 50, 100, and 200nm membrane filters, respectively. The size of the extruded liposomes was further confirmed through negative stain imaging (Inset to Fig. 1d).

#### D. Modulating fluidity of nMS ready liposomes

Fluidity of a lipid membrane can be modulated by several independent factors which include chain lengths of the lipids used and the percentage of cholesterol in the lipid membrane. In this current work, for Extended Data Fig. 5, to make liposomes with varying fluidity, we systematically varied the CH% (as reported in the Extended Data Fig. 5) in the bilayer. The change in membrane fluidity was measured by FRAP experiments on the lipid mixtures used to form the liposomes.

### MASS SPECTROMETRY ANALYSIS

#### A. Native MS analysis in Q Exactive UHMR (Thermo Fisher Scientific)

All proteins were first checked for exact mass measurement without incorporations into lipid vesicles. For this purpose, all proteins were buffer exchanged to 200 mM ammonium acetate, 2 mM DTT with Zeba^™^ spin desalting columns (Thermo Fisher Scientific). The protein concentration was kept in the range between 2μM and 10μM. Stable electrospray ionization was achieved using in-house nano-emitter capillaries. The nano-emitter capillaries were formed by pulling borosilicate glass capillaries (O.D – 1.2mm, I.D – 0.69mm, length – 10cm, Sutter Instruments) using a Flaming/Brown micropipette puller (Model P-1000, Sutter Instruments). After the tips were formed using this puller, the nano-emitters were coated with gold using rotary pumped coater Q150R Plus (Quorum Technologies). These nano-emitters were used for all nativeMS analyses. All nativeMS analysis were performed in Q Exactive UHMR (Thermo Fisher Scientific). To perform nativeMS of proteins from lipid vesicles, the nano-emitter capillary was filled with the prepared proteo-lipid vesicles and installed into the Nanospray Felx^™^ ion source (Thermo Fisher Scientific). The MS parameters were optimized for each sample. The parameters are as follows: spray voltage was in the range between 1.2 – 1.5 kV, the capillary temperature was 270^0^C, the resolving power of the MS was in the range between 3,125 – 50,000 (for MS analysis) and 100,000 (for MS/MS analysis), the ultrahigh vacuum pressure was in the range of 5.51e-10 to 6.68e-10 mbar, the in-source trapping range was between 50V and 300V. The HCD voltage was optimized for each sample ranging between 0 to 200V. We observed that for certain proteoliposomes the use of a small amount of supercharger reduced the energetic requirements to ablate the IMPs form the bilayer and enhanced the detection of membrane proteins out of lipid bilayer. Accordingly, Glycerol 1,2-Carbonate (GC) was used while studying semiSWEET, MelB, LacY, MscL without GFP tag, and AqpZ from the bilayer. For the MS/MS analysis of the lipid-bound VAMP2, the 4+ charge state was isolated in quadrupole and fragmented in the HCD cell. The isolation of the lipid-bound peak was achieved just by applying in-source trapping voltage without any HCD energy. During MS/MS, the HCD energy was set to 100V. Nitrogen was used as collisional gas. All the mass spectra were visualized and analyzed with the Xcalibur software and assembled into figures using Adobe illustrator.

For semiSWEET oligomer percentage calculation, we used two alternative approaches. In the first approach, we used UniDec^54^ (Fig. 2f and Extended Data Fig. 10). The outcome of spectra deconvolution in UniDec is shown in the inset of respective spectra in Extended Data Fig. 10 and in Fig. 2f. After spectral deconvolution, the intensity of monomer mass, dimer mass along with lipid-bound monomer and dimer mass were taken for calculations of the percentage of oligomers in detergent, 0%CL, 2.5% CL, and 10% CL containing bilayer and the final bar graph of this analysis is shown as an inset in Fig. 2e and the final mass plot is shown in Fig. 2f and Extended Data Fig. 10. In the second approach, the individual peaks were plotted in the Origin software and then the area under the curve was obtained by fitting the peak with constant baseline mode. The 6+ monomer charge state and 9+ and 11+ dimer charge states were considered for oligomer % calculation. The peak areas of protein and lipid-bound protein from Origin software were then taken for percentage calculations and the final bar graph of this calculation is shown in inset in Extended Data Fig. 10a.

#### B. Lipidomics analysis in Agilent Quadrupole Time-of-Flight 6546 mass spectrometer

Extracted *E. Coli* lipids were diluted 300 times in 4:1 MeOH: CHCl3 and spiked with Avanti Splash mix standards. 3μl of this mixture was loaded onto an Agilent Poroshell 120 (EC-C18 2.7 um, 1000bar, 2.1 x 100 mM) column using an Agilent 1290 Infiniti II LC. Mobile Phase A (60% Acetonitrile, 40% H20, 7.5 mM Ammonium Acetate) and mobile phase B (90% IPA, 10% Acetonitrile, 7.5 mM Ammonium Acetate) were used to separate peaks for detection on an Agilent Quadrupole Time-Of-Flight 6546 mass spectrometer. The gradient started at 85% (15%B) and decreased to 70% A over 2 minutes and then 52% A over 30 seconds. The gradient then slowly decreased to 18% A over 12.5 minutes and then 1% A in one minute and these concentrations held for four minutes. The gradient is then restored to 85% A and the column is washed for 5 minutes. Lipids were identified using MSDIAL 4.7^55^. Further, these identified were manually validated through observation of class specific fragment ions and class specific retention time window. The molar concentrations were normalized to the amount of Avanti Splash mix standards spiked onto each of the samples.

### BULK FUSION ASSAY

Bulk fusion assay was performed as previously described ^36^. Briefly, for the fusion assay 1 mol% NBD-PE and 1 mol% Rhodamine-PE were used along with other lipids in the v-SNARE (VAMP2) containing vesicles. The t-SNARE vesicles were unlabeled. During the fusion assay, NBD is excited at 460 nm and its emission is read at 538 nm. The Rhodamine-PE excites at 560 nm and emits at 583 nm. Since in the v-SNARE vesicles the fluorescent lipids NBD-PE and Rhodamine-PE are in close proximity, they quench each other. Consequently, upon excitation of 460nm, the NBD emission signal at 538 nm was not detected from the V-SNARE vesicles. During bulk fusion assay, prewarmed at 37°C non-labeled t-SNARE vesicles (45 μl) was mixed with labeled v-SNARE vesicles (5 μl) and the fluorescence reading was recorded immediately. Because of the fusion of v-SNARE with t-SNARE, the NBD-PE and Rhodamine-PE become distant and emission signal at 538 nm starts emerging. This signal is continuously recorded at a 30-second interval. After 90 minutes of recording, 2.5 %(w/v) DDM was added to each well and again the plate was read for 5 minutes to get the maximum emission signal of NBD-PE and the data were normalized as previously described^36^. The fusion was plotted as the percentage of the NBD-PE fluorescence. For the final comparison between different conditions, the fusion at the 21^st^ minute was taken. In all experiments, as a control, we incubated the t-SNARE vesicles (45μl) with the excess soluble cytoplasmic domain of VAMP2 (CDV) (5μl) that competes with the full-length VAMP2 in v-SNARE vesicles for binding the t-SNARE and consequently quenches the vesicle fusion.

### NEGATIVE STAIN EM PROTOCOL

Lipid vesicles were diluted 1:150 in 50mM Tris-HCl pH7.4. Five microliters of sample were applied to a carbon, Type-B 400 mesh, Cu grid that was previously glow discharged for 30 seconds. The sample was kept on the grid for 1 minute then removed with blotting paper. Five microliters of 2% uranyl formate (UFo) were applied to the grid for 5 seconds then removed with blotting paper. Immediately, 5μl UFo was applied to the grid and kept for 1 minute. The stain was removed with blotting paper, and the grid was left to dry for 30 minutes. All images were acquired using a JEOL JEM 1400-plus 120kV.

### MOLECULAR DYNAMICS SIMULATION

For the molecular dynamics (MD) simulations of VAMP2, we used the solution NMR structure, PDB ID: 2KOG^56^, taking only the ordered region (residues 26-116), and using structure 1 of the NMR ensemble. Martinize (v2.6) was used to convert the atomistic NMR structure to a coarse-grain (CG) MARTINI model, using the Martini 2.2 forcefield^57,58^ and ElNeDyn elastic network^59^.

The INSANE python script^60^ was used to generate a bilayer and embed the CG protein structure therein. The lipid composition of this membrane was selected based on the Takamori *et al*.^38^, to ensure that the simulations were performed in a similar membrane environment to that of synaptic vesicles *in vivo*. Thus, the membrane was comprised of di-C16:3-C18:3 phosphatidylethanolamine (PE), di-C16:3-C18:3 phosphatidylserine (PS), cholesterol (CH), and C16:0-18:1 phosphatidylcholine (PC), in the ratio 25:10:45:20. The box size was 17nm by x 17nm x 15nm and a physiological salt concentration of 0.15M was used. Three repeats were carried out, each of 10ms.

All simulations were performed using GROMACS 2019.1^61,62^. A time step of 20 fs was used and periodic boundary conditions were applied. The temperature was maintained at 323 K using a V-rescale thermostat, and the pressure was maintained at 1 bar using a Parinello-Rahman barostat^63^, with a time coupling constant of 12 ps and compressibility of 5 x 10^-6^ bar^-1^. Lennard Jones and Coulomb interactions were cutoff at 1.1 nm and potential modifiers were used^64^. Simulation analysis was performed using the GROMACS 2019.1 radial distribution function and plots prepared using Xmgrace^62^. The RDF was calculated for residues 26-87 of VAMP2.

### FLUROSCENCE RECOVERY AFTER PHOTOBLEACHING (FRAP) EXPERIMENTS

FRAP experiment on supported bilayer was performed as previously reported^65^. Briefly, supported lipid bilayers were formed onto the glass-bottom part of a petri dish using the Langmuir-Blodgett deposition technique^66^. Glass-bottom petri dishes (35 mm dishes from MatTek with uncoated glass cover slip of 14 mm diameter, thickness number 1.5) were soaked for 1 hour at ~ 60°C in 2% v/v cleaning detergent (MICRO-90, VWR), thoroughly rinsed with 18.2 MΩ ultra-pure water, and then dried under a stream of nitrogen. A chloroform solution of DOPC constituting the inner monolayer was first spread on degassed water in a NIMA Langmuir trough (model 611 equipped with the PS4 surface pressure sensor) and allowed to dry for 15 min at room temperature. After solvent evaporation, the film was compressed up to 33 mN/m and the monolayer was transferred onto the glass cover slip (with its lipid headgroups facing the glass surface) as the petri dish was slowly (0.5 cm/min) raised out of water. This first monolayer was allowed to dry for at least 30 min at room temperature. Meanwhile, a chloroform solution of the lipid mixture constituting the outer monolayer was spread at the air/water interface. This lipid mixture is a series of increasing cholesterol mol% (0, 15, 25, and 35) with other synaptic vesicle lipids and 1mol% NBD-DOPE. After 15 minutes, this second monolayer was transferred at a constant pressure of 33 mN/m and with a dipping speed of 0.5 cm/min onto the first monolayer (with its lipid headgroups facing the aqueous medium). After this second deposition, the bilayers remained immersed in aqueous medium throughout experiment.

FRAP experiments were performed on a confocal microscope TCS SP8 from Leica equipped with the LCS software, using an HC PL FLUOTAR 10X Dry objective (numerical aperture: 0.3; zoom: 10X) with a pinhole opened at 70.8 μm and an acquisition rate of 1 picture every 635 ms. Fluorescence bleaching and recovery were conducted as follows. For NBD: λ_exc_ = 488 nm; λ_em_ = 500–600 nm with 1 scan at 100% laser power for bleaching, and monitoring recovery at 2% of the maximum laser power. For each sample, recovery curves (average over 3 independent experiments, *i.e*. performed on a different region of the sample using the same bleaching conditions) were fitted with the software Mathematica (FRAP experiments described by modified Bessel functions; code provided upon request). In order to take into account the contribution of fluorescence recovery that occurs during the photobleaching phase and can thus affect data analysis^67^, the time t = 0 of the recovery phase in all FRAP experiments was set as the time of the last bleaching frame. When working with photosensitive probes, such as NBD, one has to correct for the intrinsic photobleaching that occurs during the recovery phase and can also affect data analysis^68,69^. In FRAP experiments, we measured the intrinsic photobleaching in a region far away from the bleaching zone. The corrected fluorescence recovery signal was then calculated by dividing the raw fluorescence recovery signal by the intrinsic photobleaching signal. Diffusion coefficient (D) of 18:1 NBD-PE lipids in the outer leaflet of different supported lipid bilayer containing different amount of cholesterol were deduced by varying the area of the bleached region (disks of diameters d= 5, 10, 15 μm). The linear relationship between the bleaching area d^2^ and the recovery time (τ) proves that lipid diffusion is controlled by Brownian motion and then diffusion coefficient (D) is calculated from the slope of these straight lines: D = d^2^/16τ where D, the diffusion coefficient is in μm^2^/s, d is bleached regions diameter, and τ is the recovery time in [s]. The diffusion coefficient (D) is then plotted against the mol% of cholesterol in the bilayer. As can be seen from the plot (Extended Data Fig. 5b), with increase in mol% of cholesterol in the bilayer the diffusion coefficient decreases. That means the fluidity of the bilayer decreases with increase in cholesterol in the bilayer.

---

**Extended Data Fig. 1:**
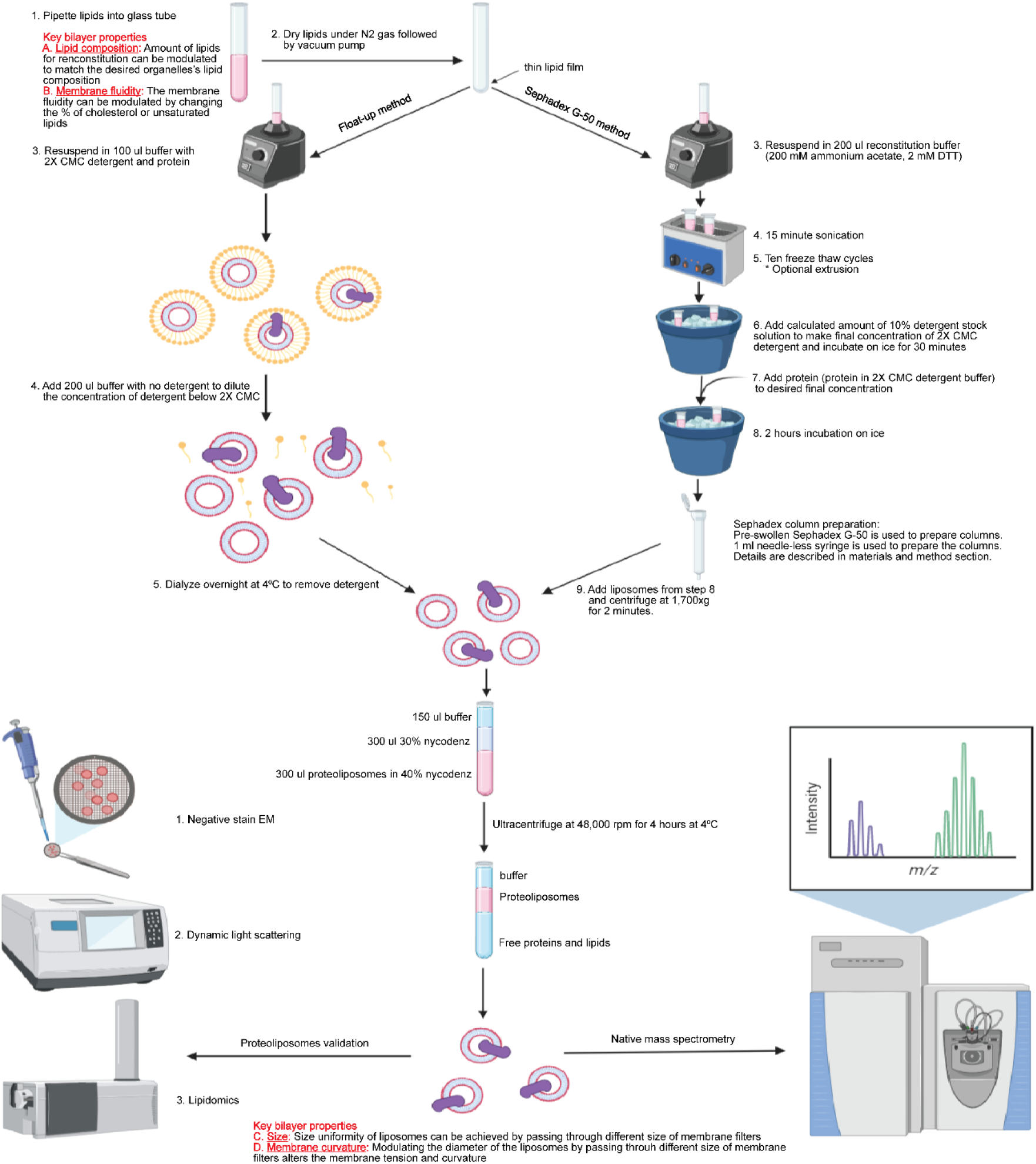
Detail flow chart of two alternate approaches that can yield native-MS ready proteoliposomes with desired lipid composition and membrane biophysical properties. The formed proteoliposomes can be isolated from the free protein and the empty liposomes through a floatation assay. For better visualization, we typically add 1% RhodPE in our lipid mix. Post-floatation, the quality of the liposomes are assessed via negative stain imaging and DLS. An optional step can be introduced to control the diameter, and in turn the curvature, of the liposomes by passing them through an extruder fitted with filters of desired dimension. The size is further assessed by negative stain or DLS. Between the two protocols, while float-up method is typically longer, for certain protein-detergent scenario, dialysis can be more effective way to reconstitute. Figure was created using Biorender.com

**Extended Data Fig. 2:**
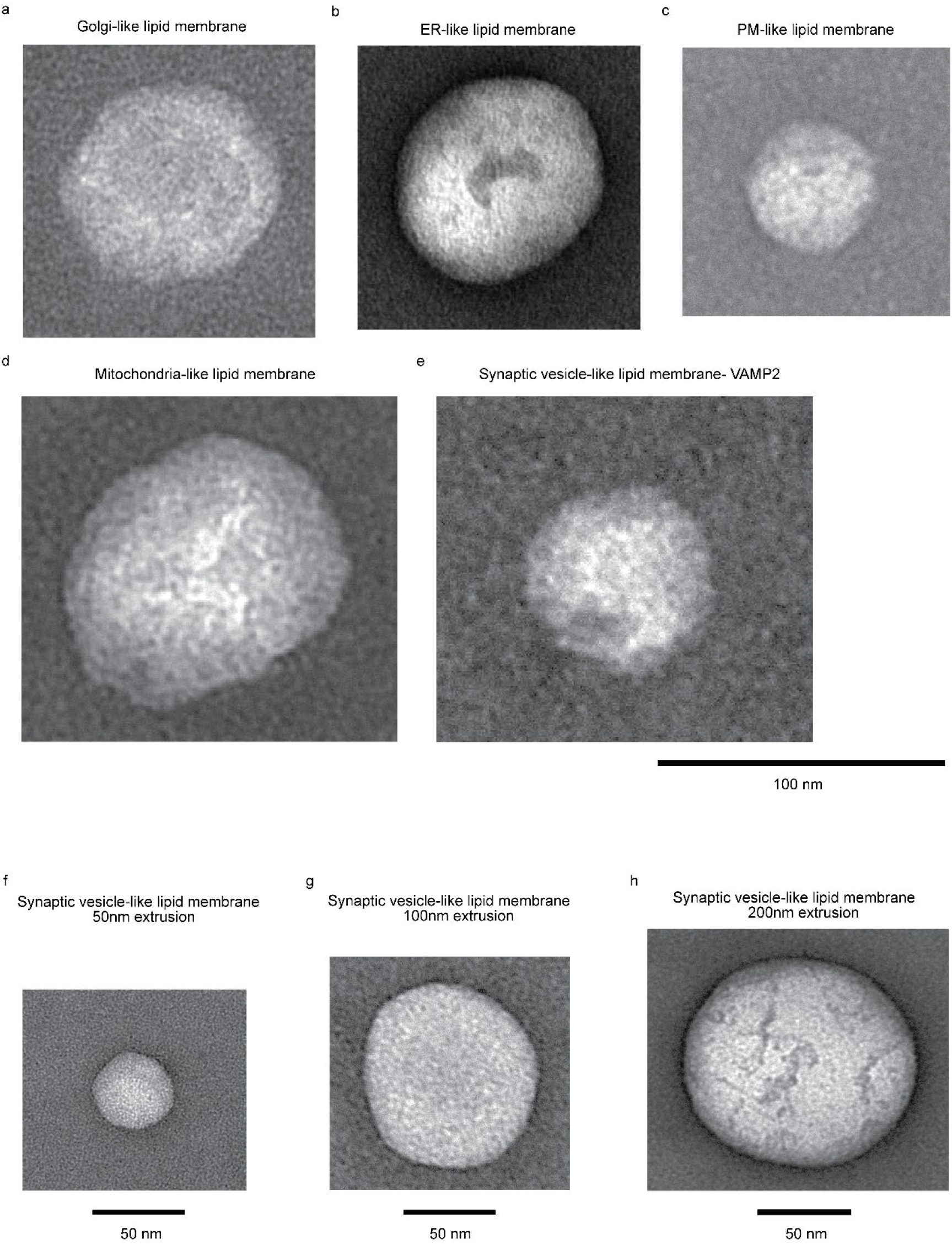
a,b,c,d, and e - Negative stain TEM images of different lipid vesicles which mimics of different eukaryotic organellar membranes. f, g, and h - Negative stain TEM images of different curvature synaptic vesicles. The desired curvature was achieved by passing the synaptic vesicles like lipid vesicles through different pore size membrane. All the sample was taken from a pool of vesicles that were subjected to nativeMS (Fig. 1), confirming the vesicles in the samples. All measurements were replicated over three independent measurements.

**Extended Data Fig. 3:**
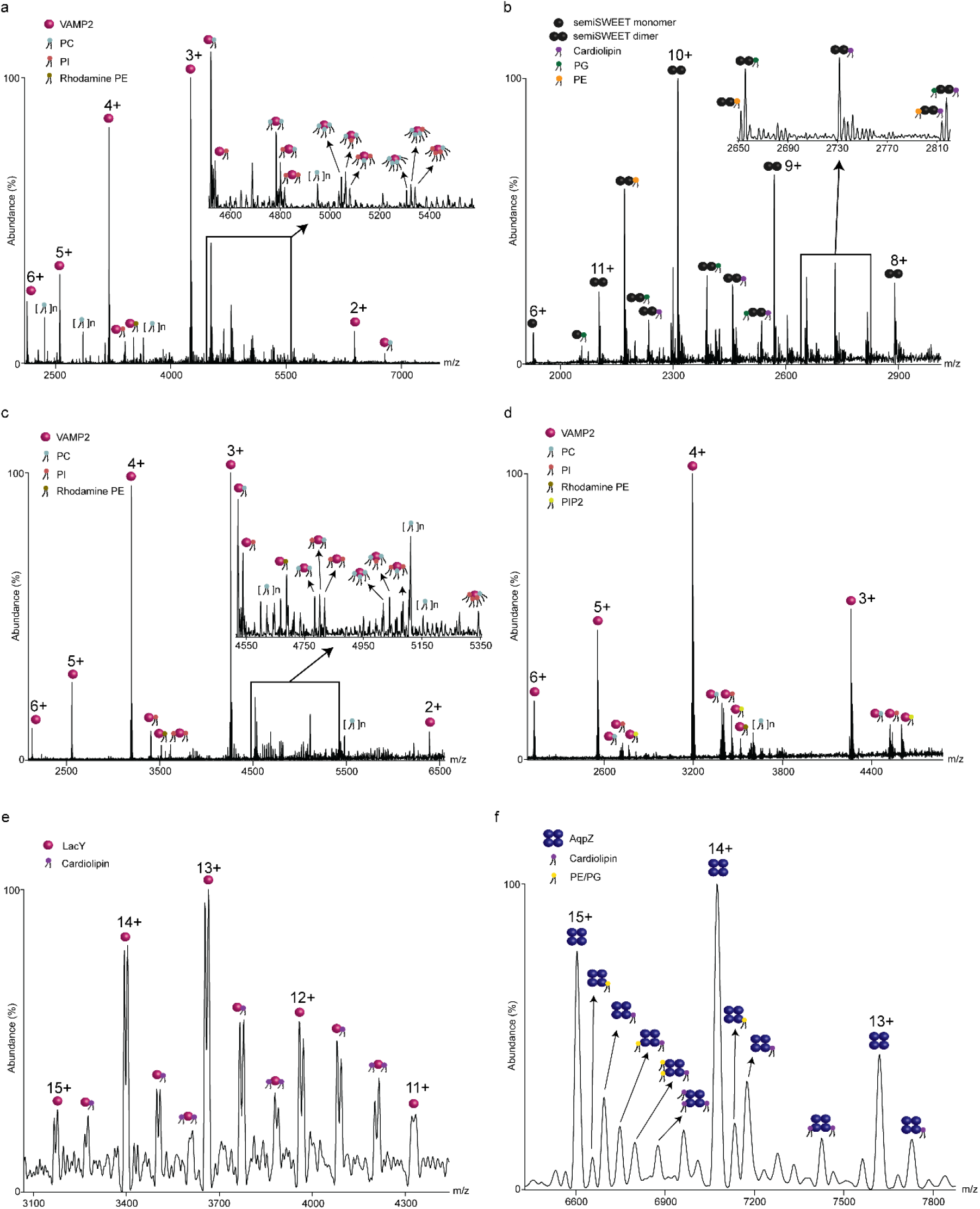
Detail assignments of the bound lipids in the spectra of a, Golgi like lipid membrane, b, mitochondria like membrane, c, ER like lipid membrane, d, PM like lipid membrane, e, and f, bacterial inner plasma membrane like lipid membrane as observed in the Fig. 1. The identity of the lipids could be determined by directly comparing the adduct masses against the theoretical masses of the individual lipids used to make the bilayer (Supplementary Table 2). All measurements were replicated over three independent measurements.

**Extended Data Fig. 4:**
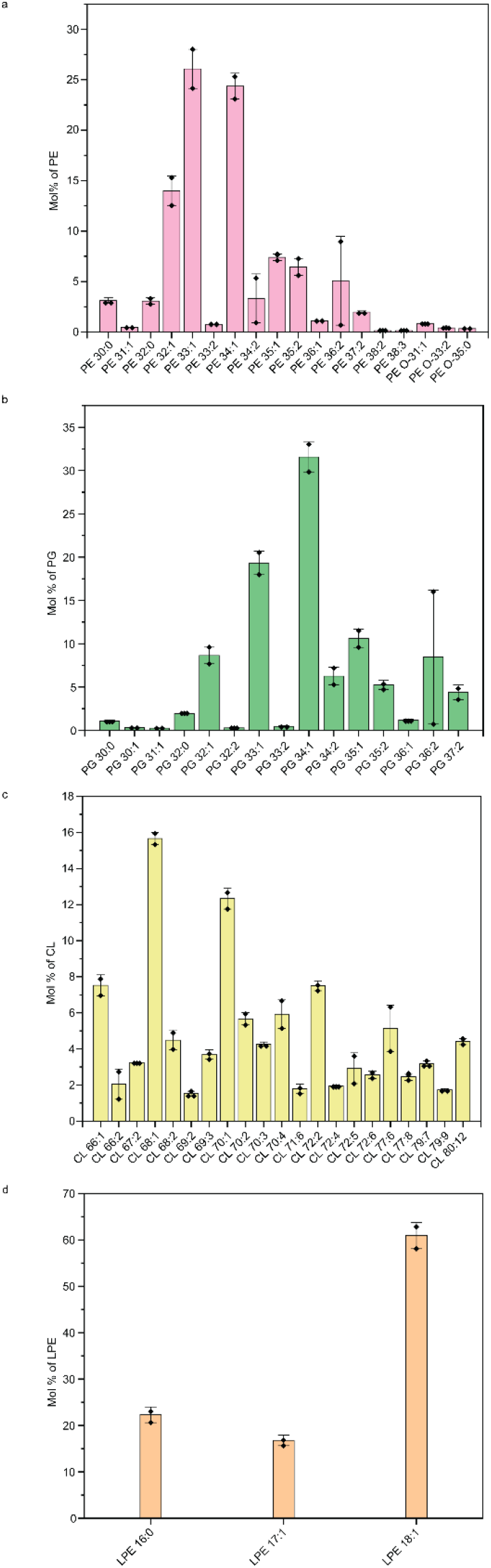
Detailed lipidomics profile of the extracted E.coli membrane used to reconstitute LacY and AqpZ (Fig. 1). For each class of lipids, the components were divided as per the total chain length distribution and quantified. The data presented as mean ± s.e. (N=2).

**Extended Data Fig. 5:**
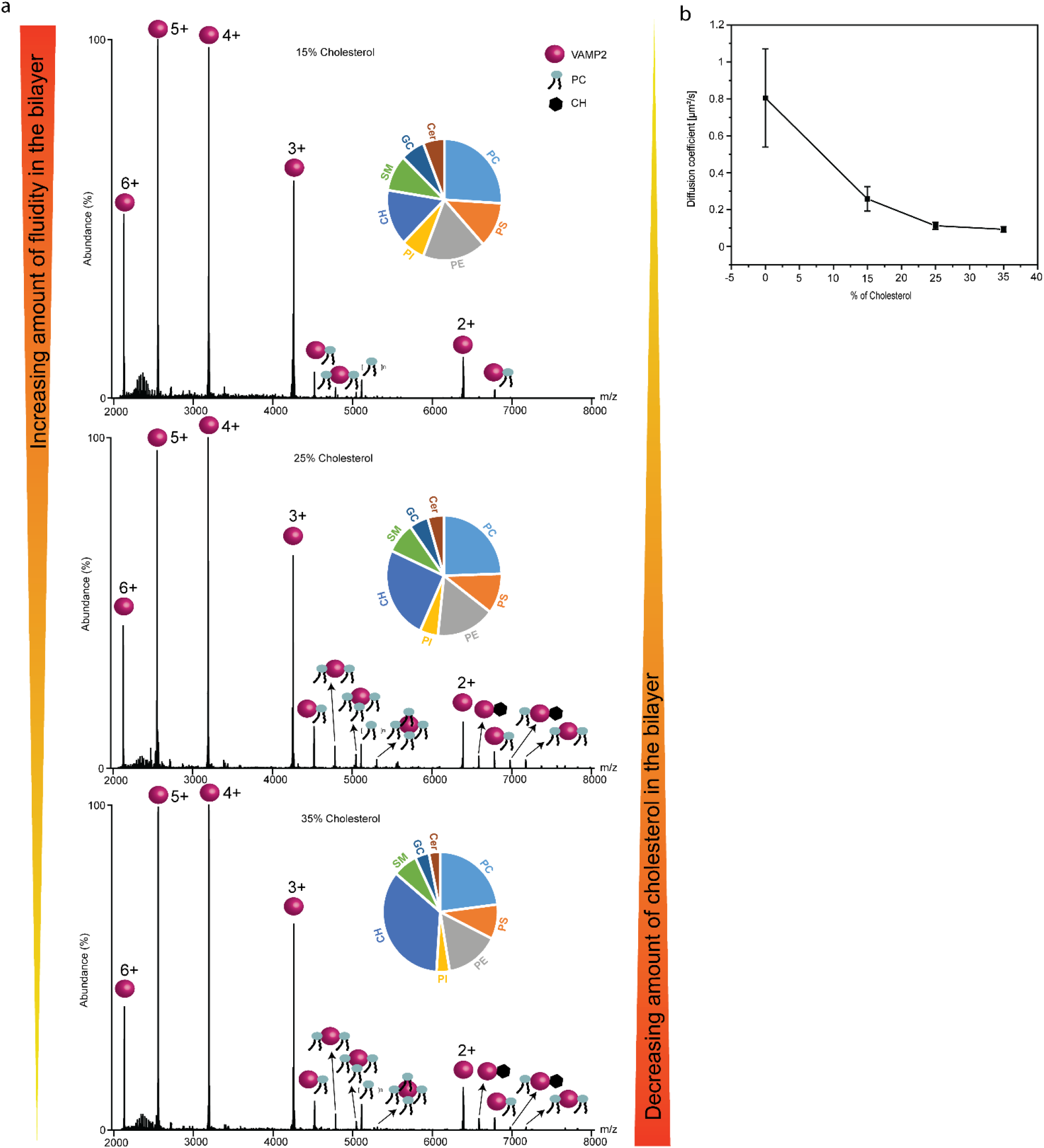
a, nMS analysis of IMP (VAMP2) from vesicles with increasing amount of cholesterol. All measurements were replicated over three independent measurements, b, The diffusion coefficient (μm2/s) plotted against the mol% of cholesterol in the bilayer. As can be seen from the plot, with increase in mol% of cholesterol in the bilayer the diffusion coefficient decreases. That means the fluidity of the bilayer decreases with increase in cholesterol in the bilayer. Each data point represents the average diffusion coefficient from three independent experiment. The data presented as mean ± s.e. (N=3).

**Extended Data Fig. 6:**
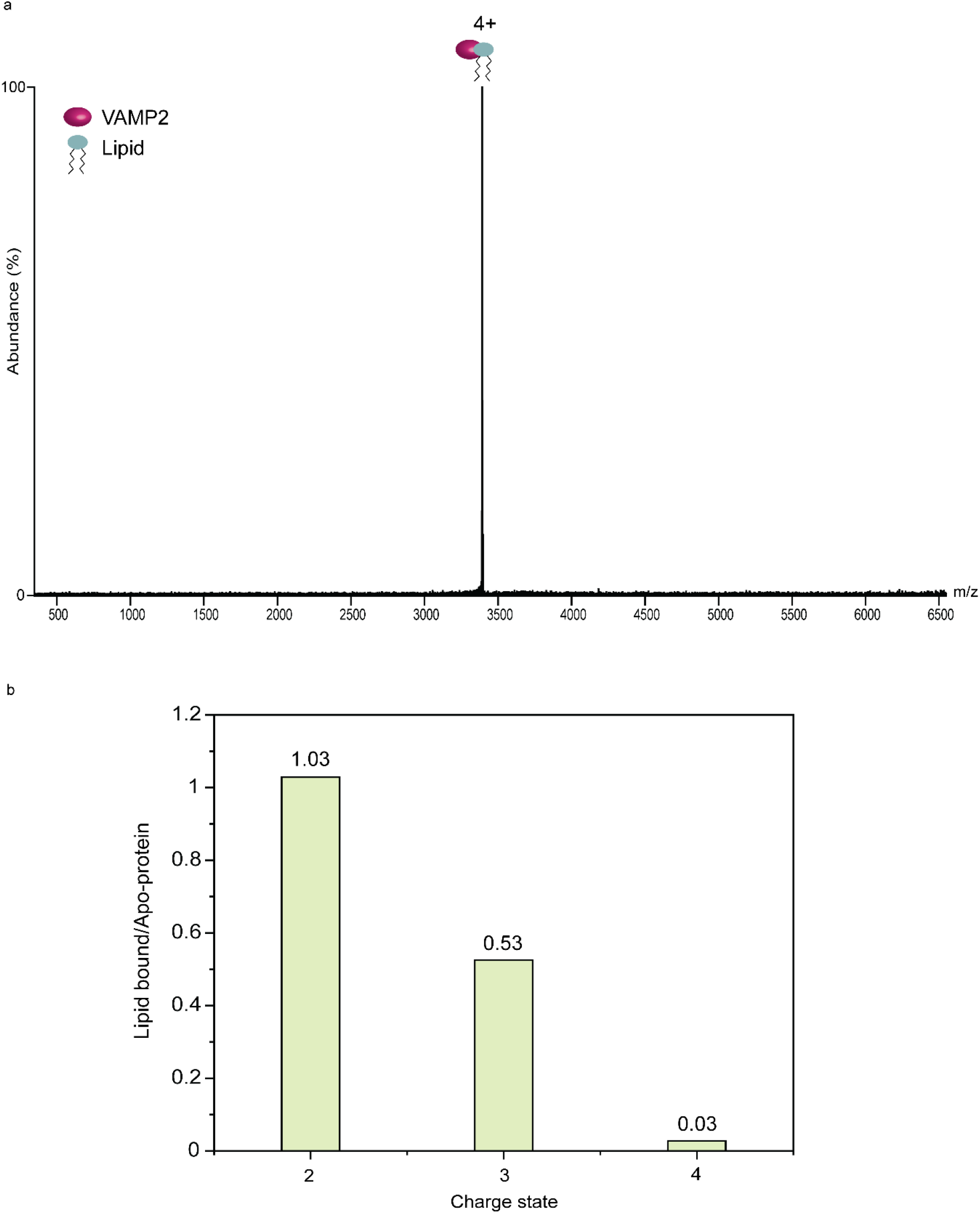
a, Isolation spectra of 4+ charge state lipid bound VAMP2 from synaptic vesicles mimicking vesicles. b, Lipid bound/Apo-protein ratio calculated from area under the curve of apo-protein peaks and lipid bound protein peaks (and then total lipid bound area taken to get ratio) in nMS spectra of VAMP2 from synaptic vesicles mimicking vesicles.

**Extended Data Fig. 7:**
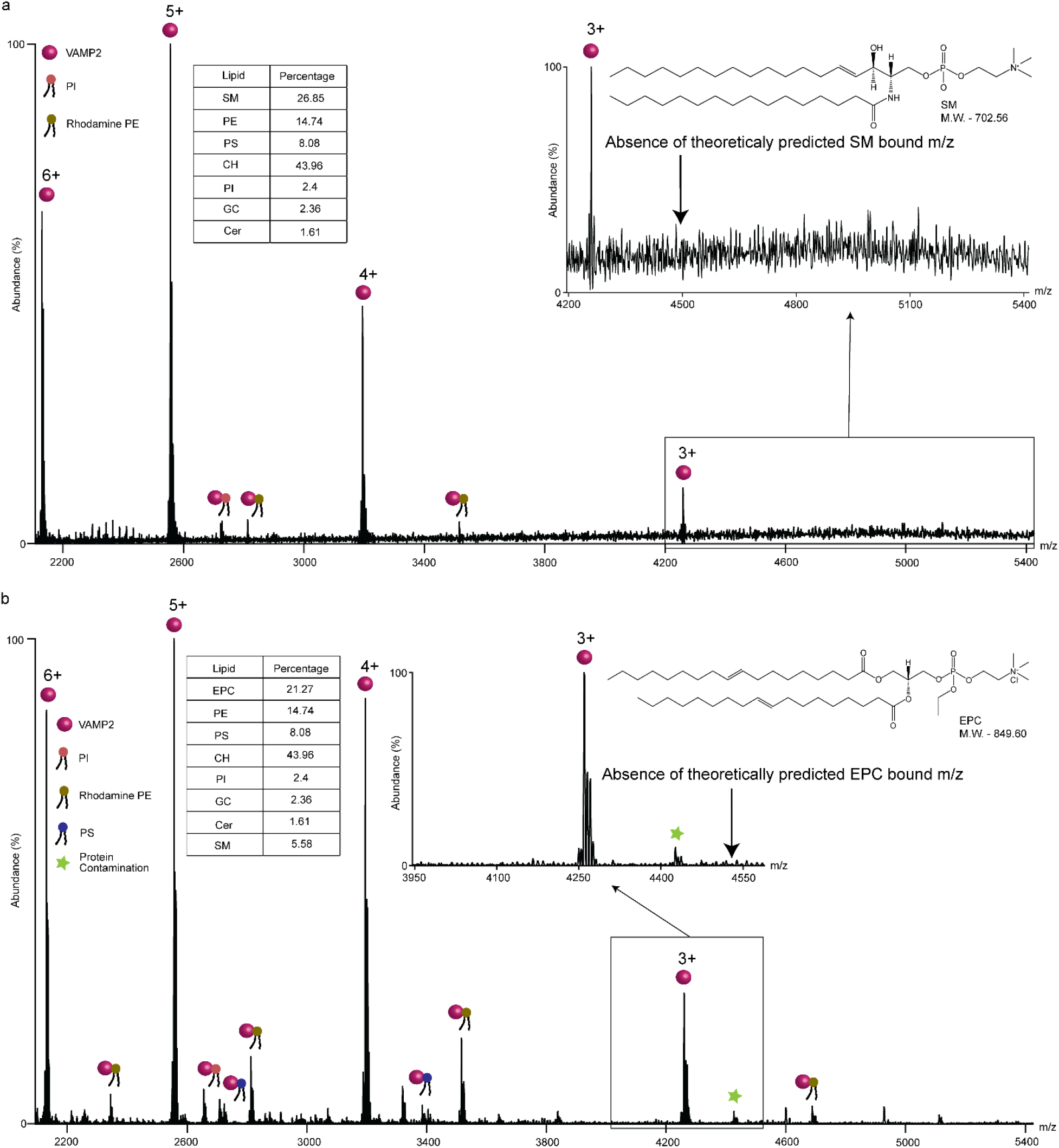
Spectra of VAMP2 from SV-like lipid vesicles where all the PC were replaced by a, SM and b, EPC (lipid compositions shown in the respective inset tables). The arrows showing the absence of theoretically predicted SM/EPC bound m/z values. In both the cases, under the same MS conditions, no detectable amount of SM or EPC binding was observed. All measurements were replicated over three independent measurements.

**Extended Data Fig. 8:**
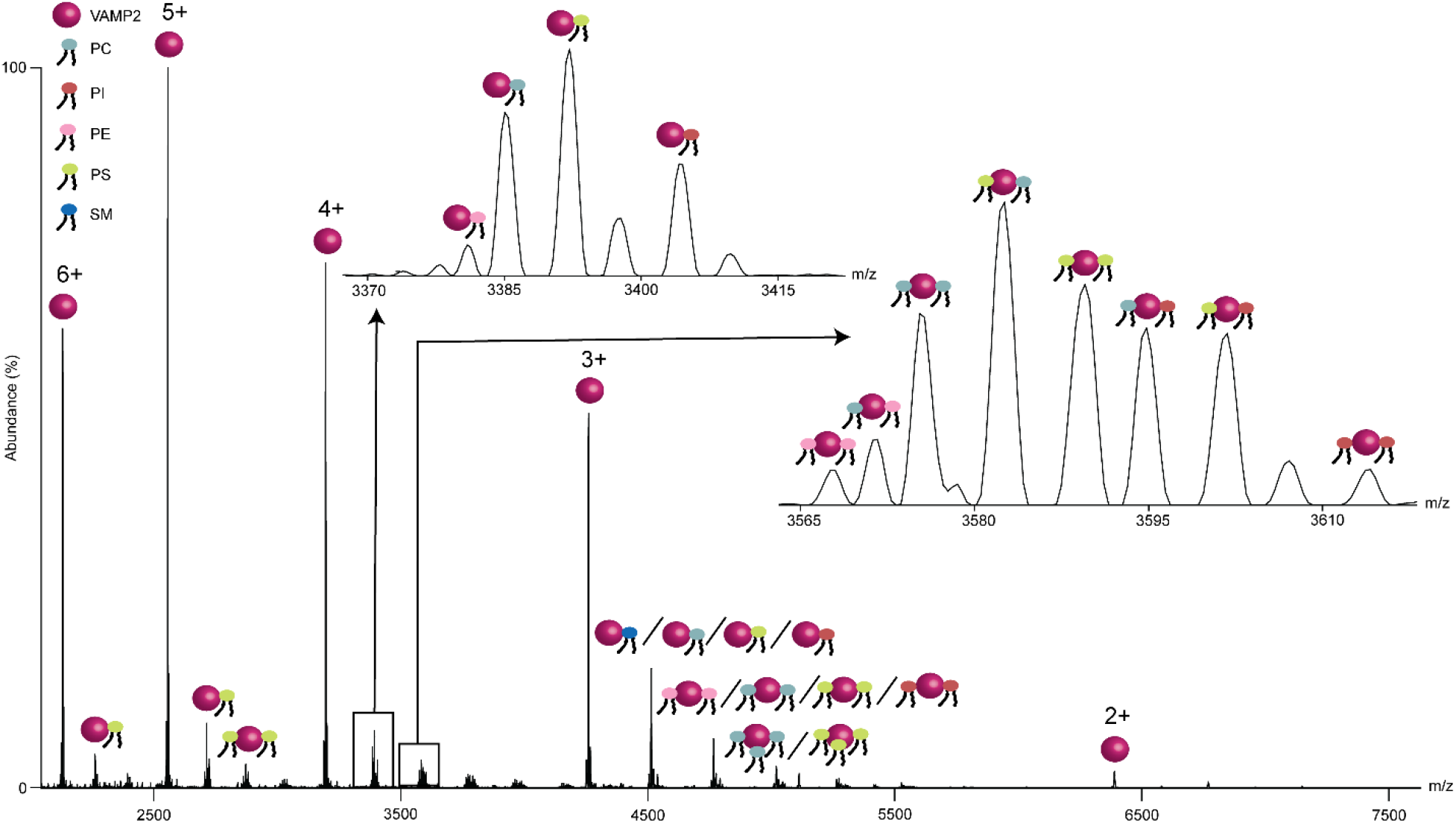
Spectra of detergent solubilized VAMP2 incubated with the same mixture of lipids present in SV-like liposomes. Taking the example of +4 charge state, the data clearly shows that VAMP2 binds to several phospholipids without any specificity to any specific lipids. Further no detectable amount of cholesterol binding was observed. The data was obtained in MS conditions identical to that used for detecting VAMP 2 from SV like liposomes. All measurements were replicated over three independent measurements.

**Extended Data Fig. 9:**
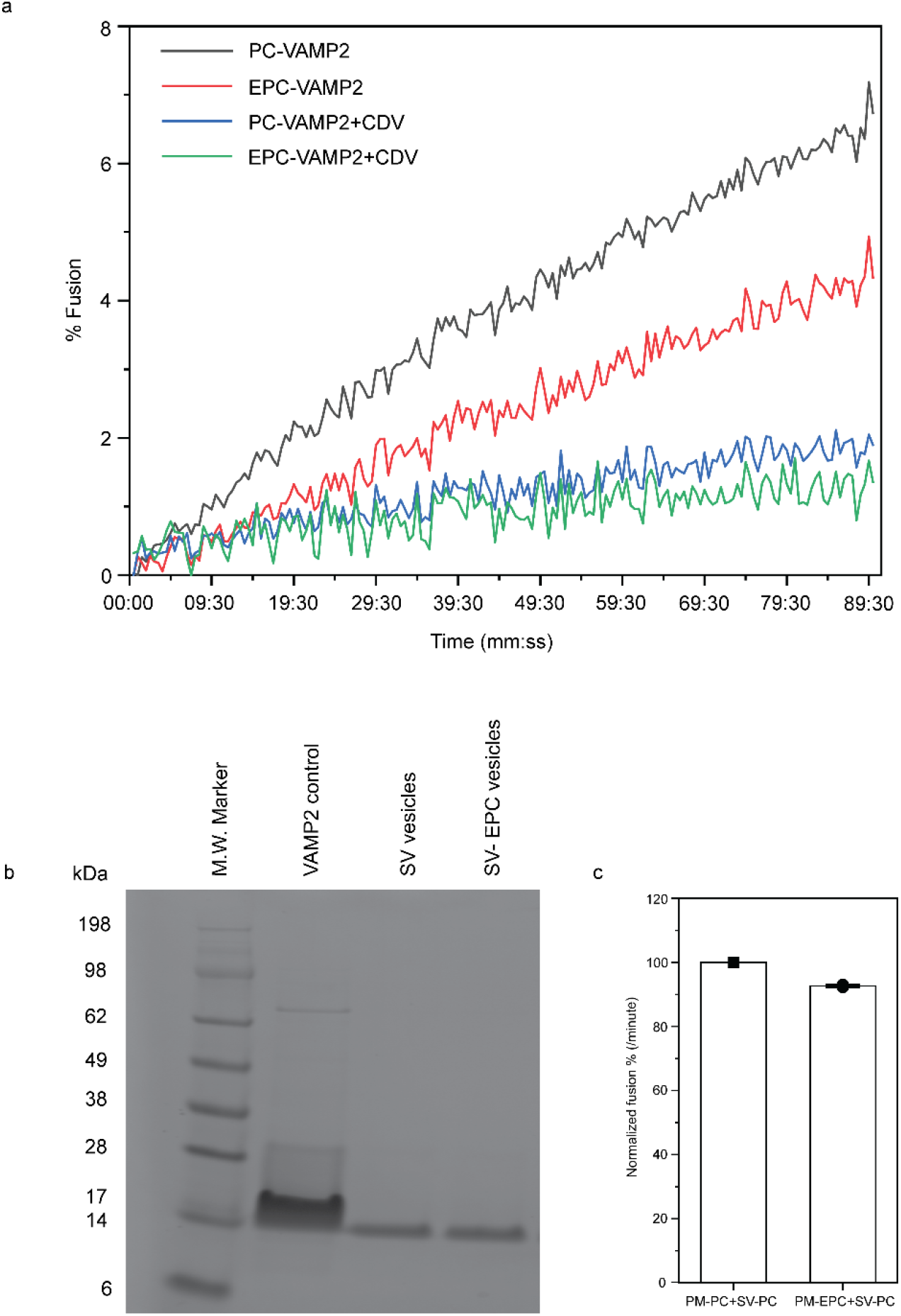
a, % Fusion vs the time course data for the VAPM2 in SV like and EPC-SV vesicles. In both the cases, CDV (cytosolic domain of VAMP2) was used a negative control. In presence of excess CDV, CDV binds and occupies the VAMP2 binding domains of the t-SNAREs. Consequently, upon addition of VAMP2 reconstituted vesicles, these vesicular VAMP2s cannot bind to t-SNARE anymore and no fusion is observed. As seen, the EPC-SV vesicles, devoid of PC, fuses with the PM at a significantly lower rate than the PC containing native-like SV liposomes. b, SDS-Gel picture showing similar band intensity of VAMP2 in SV like and EPC-SV vesicles. c, Vesicle fusion assay performed between t-SNARE in PM like lipid composition vesicles (tSNARE-PM) and VAMP2 in SV like vesicles (VAMP2-PC). Subsequently, to assess the effect of EPC on fusion rate, PC was replaced with EPC in PM like vesicles containing t-SNARE. Similar rate of fusion was observed as observed in t-SNARE in PM like vesicles with PC. This confirms that the reduced fusion rate observed for SV-EPC vesicles (Fig. 2), is not because of replacing PC with EPC in the bilayer, but due to abrogation of VAMP2-PC binding. The data presented as mean± s.e (N=5).

**Extended Data Fig. 10:**
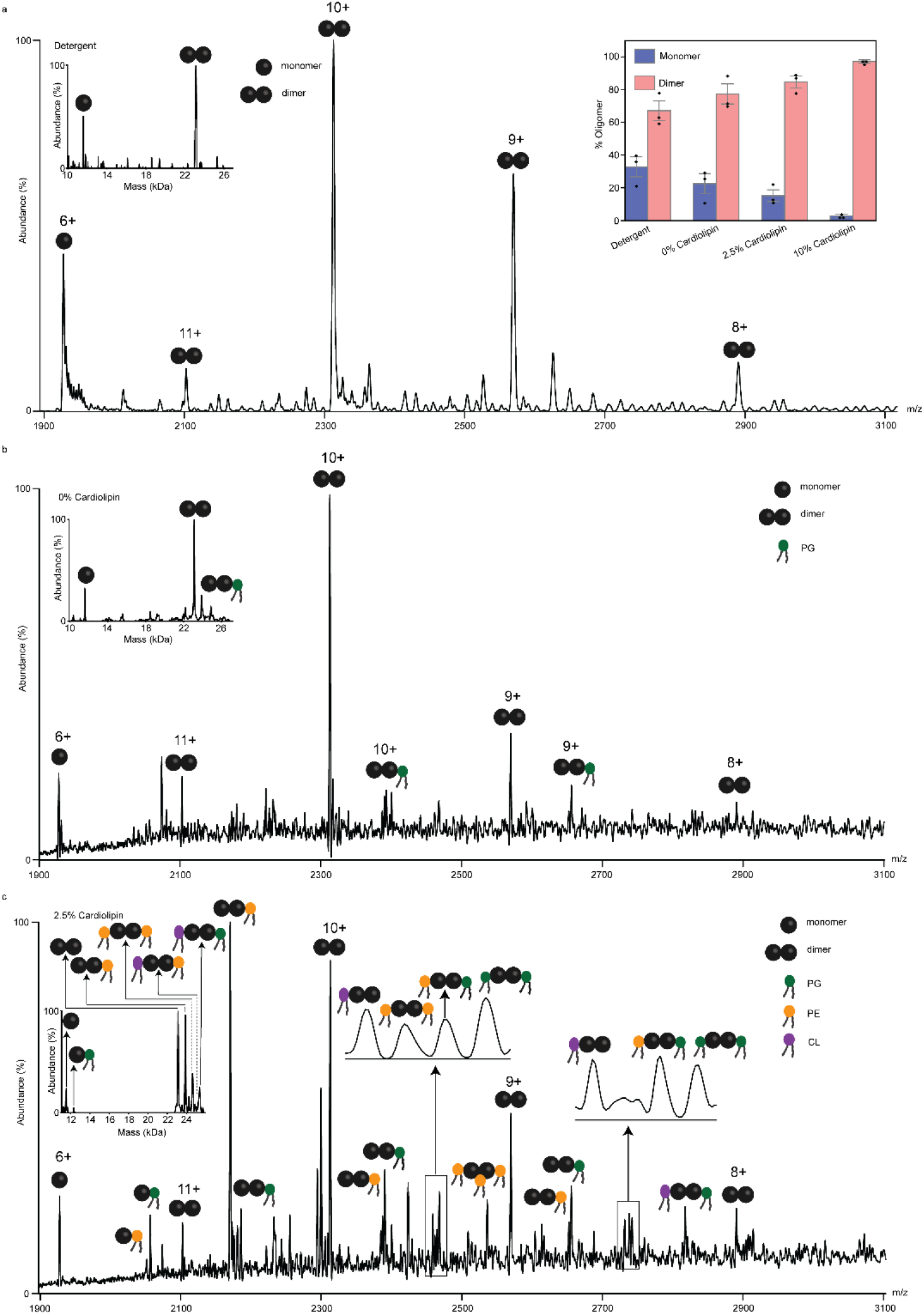
nativeMS spectra of semisweet from a, detergent, b, bilayer with PE, PG without CL, c, bilayer with PE, PG, and 2.5% CL. The individual peaks were plotted in the Origin software and then the area under the curve was obtained by fitting the peak with constant baseline mode. The 6+ monomer charge state and 9+ and 11+ dimer charge states were considered for oligomer % calculation. The peak areas of protein and lipid-bound protein from Origin software were then taken for percentage calculations and the final bar graph of this calculation is shown as an inset in a. The data clearly shows increase in CL% increases the dimeric population in the bilayer. The inset on the left side in each panel shows the mass plot obtained from UniDec analysis.

